# Sex-stratified Gut Microbiome Disruption is Associated with Altered Hepatic Gene Expression during Acute Azoxystrobin Exposure

**DOI:** 10.64898/2026.02.18.706612

**Authors:** Luoyan Duan, Wes Arend Baumgartner, Jeremiah W. Wanyama, Lydia Okyere, David Anthony Alvarado, Bushra Fazal Minhas, Christopher A Gaulke

## Abstract

Azoxystrobin is a widely used fungicide that has been associated with to reproductive, neurological, and developmental defects. This chemical also disrupts gut microbial communities; however, if these perturbations contribute to the harms associated with exposure to azoxystrobin, this remains unclear. In this study, we investigated the effects of acute exposure to a series of concentrations (5–500 mg/kg) of azoxystrobin on the host and gut microbiota in zebrafish. Fecal amplicon and shotgun metagenomic sequencing was integrated with liver gene expression to quantify associations between microbiome disruption azoxystrobin toxicity in the host. Azoxystrobin exposure resulted in significant alteration in microbiome composition and functional potential in a dose- and sex-dependent manner. Microbial communities in exposed animals exhibited an increased abundance of xenobiotic metabolism pathways and decreased bacterial motility and lipopolysaccharide biosynthesis pathway metabolism. At the host level, histopathology identified increased biliary proliferation, most evident in medium- and high-dose fish. We also observed hepatic transcriptional changes consistent with a stress response, including altered redox-associated genes and reduced expression of lipid and small-molecule metabolic genes, with sex-stratified differences. Importantly, alterations in host transcriptional programming correlated with the compositional changes in exposed microbiota. Together, these results suggest concurrent impacts of azoxystrobin on gut microbiota and the liver implicate the microbiome as a potential contributor to changes in liver gene expression during exposure.

**Importance:** Widespread fungicide use contaminates ecosystems worldwide, but the biological pathways underlying their effects on humans and other animals are not well understood. Using zebrafish (*Danio rerio*), we found that short-term exposure to the fungicide azoxystrobin was associated with changes in the gut microbiome, liver gene activity, and liver changes. Exposure produced dose- and sex-dependent shifts in microbial communities, including changes in predicted microbial functions involved in chemical metabolism, bacterial motility and defense. Compositional changes in the microbiome correlated with gene-expression changes consistent with stress and altered metabolism in exposed fish, suggesting that exposure induced disruption may contribute to exposure impact to the host. These results highlight a potential role for the microbiome in mediation of the impacts of azoxystrobin on host physiology. As such microbial based interventions could be a viable strategy to mitigate exposure impacts on health.

## Introduction

Azoxystrobin (AZO) is a widely used broad-spectrum strobilurin fungicide commonly applied to cereals, fruits, nuts, and vegetables as well as some construction materials (1, 2). Global sales of AZO have been estimated to exceed $1.2 billion in 2024, and market projections suggest continued growth, potentially reaching over $2.5 billon by 2030 (3). Concerningly, AZO or its metabolite, Azoxystrobin acid, have been detected in produce (4, 5), surface water (6), groundwater (7), indoor dust (2, 8), and pregnant women and children (9). As a result, AZO has been flagged as a priority chemical for biomonitoring in the United States (10).

Azoxystrobin inhibits mitochondrial respiration in fungi by targeting complex III of the electron transport chain, thereby disrupting ATP synthesis and inducing oxidative stress (11). Although azoxystrobin is designed to selectively target fungal cells, its toxicological effects on non-target organisms have been described. For instance, AZO exposure induces dose- and time-dependent oxidative damage in rat testicular and liver tissues (12). Studies in zebrafish have revealed similar impairments, including reduced fertilization rates, and embryonic developmental defects (13). Azoxystrobin also induces oxidative stress, apoptosis and genotoxicity in zebrafish larvae and adult livers (14, 15). Similarly in mice, AZO causes inflammation and decreased expression of key determinants of gut barrier integrity including Occludin and Zonula occludens-1 (ZO-1) (16). These findings underscore the importance of continued toxicological evaluation and environmental monitoring of AZO to better understand its potential risks to non-target organisms and human health.

The gut microbiome plays a vital role in maintaining host health, contributing to essential processes such as nutrient absorption and immune system regulation (17). Growing evidence also links microbiome to metabolism of xenobiotics, including environmental contaminants such as pesticides (17, 18). Gut microbes encode a suite of enzymes with the potential to detoxify xenobiotics, including β-glucuronidases, esterases, oxidoreductases, and transferase (18). For instance, *Lactobacillus plantarum* strains have been shown to degrade organophosphorus compounds such as dimethoate, phorate, omethoate, and chlorpyrifos via esterase-mediated hydrolysis (19). Conversely, microbes can also enhance the toxicity and bioavailability of xenobiotics or their metabolites through several mechanisms (20) including reactivation of detoxified metabolites by β-glucuronidase–mediated deconjugation which can exacerbate gastrointestinal toxicity (21). Pesticide exposure also impacts microbial communities, shifting microbiome composition and interactions with their host (22). For example, Motta *et al.* demonstrated that glyphosate exposure at field-relevant doses significantly perturbs the gut microbiota of honeybees. They found that glyphosate impaired the beneficial bacterium *Snodgrassella alvi*, which led to a weakened resistance to opportunistic pathogens and increased bee mortality (23). Similarly, adverse effects of AZO in mice, including colonic inflammation and impaired epithelial barrier integrity, were found to be mediated by alterations in the gut microbiota (16). However, despite growing evidence that pesticides can reshape the gut microbiome, it remains unclear whether and how these microbiome disruptions contribute to host-level toxicity, particularly for widely used fungicides such as AZO.

To better understand the potential role of the microbiome in AZO responses, we examined how acute AZO exposure affects gut microbiome composition and inferred functional potential in a zebrafish model. Zebrafish were continuously fed AZO-contaminated food for 14 days. Microbial community profiling was assessed via 16S rRNA sequencing, and functional potential was quantified through shotgun metagenomics. By integrating these data with liver transcriptomics, we identify potential links between AZO disruption of the microbiome and pesticide toxicity. Our findings offer new insights into the impacts of AZO on microbial communities and how these effects may alter response to toxicants.

## Materials and Methods

### Animal Husbandry

Adult tropical 5D (T5D) zebrafish were housed in a flow-through aquatic system (Aquaneering, San Marcos, CA) with automatic 50% water exchange every 6 h using conditioned, heated (28°C) reverse osmosis water. Water quality was maintained 7.0-7.5 pH, 500 −1200 μS, 0 ppm NH_3_, 0 ppm NO_2_^-^, 0 ppm NO_3_^-^. Fish were kept at 28°C on a 14:10 h light: dark cycle under the protocol from the Institutional Animal Care and Use Committee at the University of Illinois at Urbana-Champaign (permit number 21188).

### Animal Exposures

Sixty (60) six-month-old male and female T5D wild-type zebrafish were randomized into four groups (11 males, 4 females per group), individually housed (1.8 L), and acclimated for 14 days on a commercial diet (Gemma Micro 300; Skretting, Westbrook, ME USA).

Azoxystrobin or vehicle dimethyl sulfoxide (DMSO; Sigma-Aldrich, Saint Louis, MO, USA) was applied to the diet using a mucosal atomization device (LMA™ MAD Nasal™, MAD300, Teleflex Inc., Morrisville, NC, USA). AZO exposure was targeted at 0, 5, 50, or 500 mg AZO/kg body weight/day. Diet concentrations were not analytically verified; doses reflect nominal targets. Each fish received 10.4 mg diet/day (calculated from an average body mass of 490 mg per six-month-old zebrafish; ∼2.12% wet body weight/day). Fish were monitored throughout the experiment to ensure that all food was consumed at each feeding. Animals were exposed to these diets for fourteen days. Fecal samples were collected on day 0 (prior to exposure), day 7, and day 14 and stored at −20°C until processing. At day 14, fish were euthanized (ice water). Seven per group were immersion-fixed in 4% paraformaldehyde (PFA); liver and gut from the remaining eight were stored at −80°C.

### 16S rRNA amplicon library preparation, sequencing and data analysis

Zebrafish fecal samples were processed for microbial DNA isolation using the DNeasy PowerSoil Pro kit (QIAGEN, Germantown, MD, USA) with a modified ten-minute incubation at 65°C to facilitate cellular lysis using TissueLyser III (QIAGEN). The bacterial 16S rRNA V4 region was amplified using the universal barcoded primers 515F and 806R (24, 25). Triplicate 25 µL reactions containing 10 µL DreamTaq (Thermo Fisher Scientific, Carlsbad, CA), 0.5 µL of each primer, 13 µL of nuclease-free water, and 1 µL of DNA template, amplified at 95 °C for 3 minutes, followed by 35 cycles of denaturation (95 °C, 45s), annealing (50 °C, 60s), and extension (72 °C, 90s), with a final extension at 72 °C for 10 minutes. Amplicons were pooled, quantified using the Qubit® HS kit (Life Technologies, Carlsbad, CA), purified with QIAquick PCR Purification Kit (QIAGEN), diluted to 10 nM and sequenced an Illumina NovaSeq (250 bp paired-end). The paired reads were then processed using DADA2 (v 1.30) for filtering (Phred < 30). Quality controlled sequence reads were denoised and amplicon sequence variants inferred with DADA2. Taxonomy was assigned using the SILVA database (v138.1) (26, 27).

Microbial community data was rarefied to 50,000 counts using the vegan package (v2.6.6.1) in R (v4.3.1). Richness and Shannon alpha diversity was modeled using a generalized linear mixed-effects model (glmmTMB v.1.1.9.9 in R) assuming a Gaussian distribution. The model included fixed effects of group, day, and their interaction, with a random intercept for animal ID. Pairwise comparisons were assessed using estimated marginal means (R, emmeans, v.1.11.1) (28), with Benjamini–Hochberg correction. Associations between microbiome beta diversity (Jaccard dissimilarity), sampling date, group, and their interaction were quantified using Permutational Multivariate Analysis of Variance (PERMANOVA, method= “jaccard”, vegan). Dimensionality reduction of microbiome data was performed with principal coordinate analysis (PCoA) in R using cmdscale (vegan, method= “jaccard”) and visualized with ggplot2 (v 3.5.1). Differential abundance was assessed using negative binomial generalized linear mixed effects models (R, glmmTMB v.1.1.9.9) (29). For each genus, two models were constructed: a reduced model containing only intercept and random effect (id) and a full model that included group, time, sex, and the interaction between group and time or sex in addition to the random effect of animal id. An analysis of variance (ANOVA) test then determined if the full model explained significantly more variation than the reduced model. False discovery rate, was controlled using q-values (qvalue v. 2.32.0) function in R.

### Shotgun metagenomics sequencing and analysis

Shotgun metagenomic libraries were prepared with the UltraLow Input Library construction kit from Tecan (Tecan, Nänikon, Switzerland) and sequenced on a NovaSeq X Plus (150 bp paired-end). The raw sequence reads per sample ranged from 63.22M to 83.22M (average 73.35M per sample). Sequencing adapters and low-quality reads were filtered with Cutadapt (30) and reads aligning to the zebrafish reference genome (GRCz11) were removed with bowtie2 (v2.5.1) (31). Clean reads were then processed with HUMAnN 3.0 (32) (UniRef90; default parameters), renormalized to counts per million (CPM), and annotated to Kyoto Encyclopedia of Genes and Genomes (KEGG) pathways. The taxonomic abundances were estimated using Kraken2 (33) and Bracken (34). Differential abundance of KEGG pathways and microbial species between exposure groups were computed using a negative binomial generalized linear model (MASS, glm.nb in R, 7.3.60.0.1). Likelihood ratio tests (ANOVA) compared full models (predictors: group, sex, interaction) against intercept-only null models. Significance was defined as FDR < 0.20 (R p.adjust).

### Transcriptome analysis (RNA-Seq)

Total RNA was extracted from zebrafish liver using the RNeasy Lipid Tissue Mini Kit (QIAGEN). Poly-A selected mRNA was utilized to construct RNA-Seq libraries with the TruSeq™ mRNA library Prep kit (Illumina, San Diego, CA, USA), and sequencing was performed on an Illumina NovaSeq (100bp single-end). The raw reads were filtered and trimmed with cutadapt (v 2.6) (30) and high-quality reads were aligned to the *Danio Rerio* genome (GRCz11) with the STAR aligner (v 2.7.10a) (35). Differential gene expression analysis was conducted in R (v4.3.2) using the likelihood ratio test implemented in DESeq2 (v.1.42.1) (36). Gene set enrichment analysis of differentially expressed gene sets (FDR < 0.05) was performed with the gprofiler2 (v. 0.2.3) (37) using default parameters. The volcano plots and heatmaps were constructed using ggplot2 (v.3.5.1) and ComplexHeatmap (v. 2.18), respectively.

### Histology

Whole zebrafish were fixed overnight at 4°C in 4% PFA diluted in phosphate-buffered saline (PBS). The animals were bisected parasagitally, placed in cassettes and processed by standard methods (tissues were dehydrated in graded ethanol solutions (70% - 100%), cleared with xylene), and embedded in paraffin. The samples were sectioned at 5 µm thickness and stained with hematoxylin and eosin (H&E). All tissues in the slide were examined, and liver sections were evaluated for histopathologic lesions, and biliary changes were recorded semi-quantitatively in a subset of fish.

### Data availability

The raw sequence files generated during the current study are available at the NCBI Sequence Read Archive (SRA) under the project accession numbers PRJNA1415316, PRJNA1416997 and PRJNA1418170, samples SAMN54929448 - SAMN54929625, SAMN54986945 - SAMN54986972 and SAMN55038798 - SAMN55038812.

## Results

### Acute AZO exposure significantly disrupted the microbiome diversity and composition

To determine the impact of AZO on the gut microbiome, we profiled microbial communities of 178 fecal samples using 16S rRNA sequencing. A total of 57,454,091 high quality reads were generated from which 4,644 amplicon sequence variants (ASVs) and 433 distinct genera were identified. Amplicon sequence variants richness remained stable, **(Figure 1A)**. Shannon diversity increased significantly in the high group at Day 7 (P < 0.001) and in both medium and high groups at Day 14 (P < 0.01) relative to controls (**Figure 1B**). Similarly, prior to exposure no significant differences were noted in microbiome diversity across groups (PERMANOVA; R^2^ = 3.40x10^-2^, F = 6.60x10^-1^, *P* = 8.80x10^-1^). However, at seven (R^2^ = 1.66x10^-1^, F = 3.89, *P* = 2.00x10^-4^) and 14 days of exposure (R^2^ = 1.28x10^-1^, F = 2.76, *P* = 1.80x10^-4^) microbiome diversity was significantly associated with exposure group (**Figure 1C-E**). The most dominant genera in the zebrafish gut microbiome included *Aeromonas*, *Cetobacterium*, *Mycoplasma*, and *Pseudomonas* (**Figure S1**). We found that the abundance of 25 genera was significantly altered in the exposed groups compared to the control group (**Table S1**; FDR < 0.2). Specifically, on Day 7, *Aeromonas* (high group) and *Shewanella* (medium and high groups) decreased significantly, while *Pseudomonas*, *Bosea*, *Defluviimonas*, and *Mycoplana* increased in the high group.

**Figure 1.**
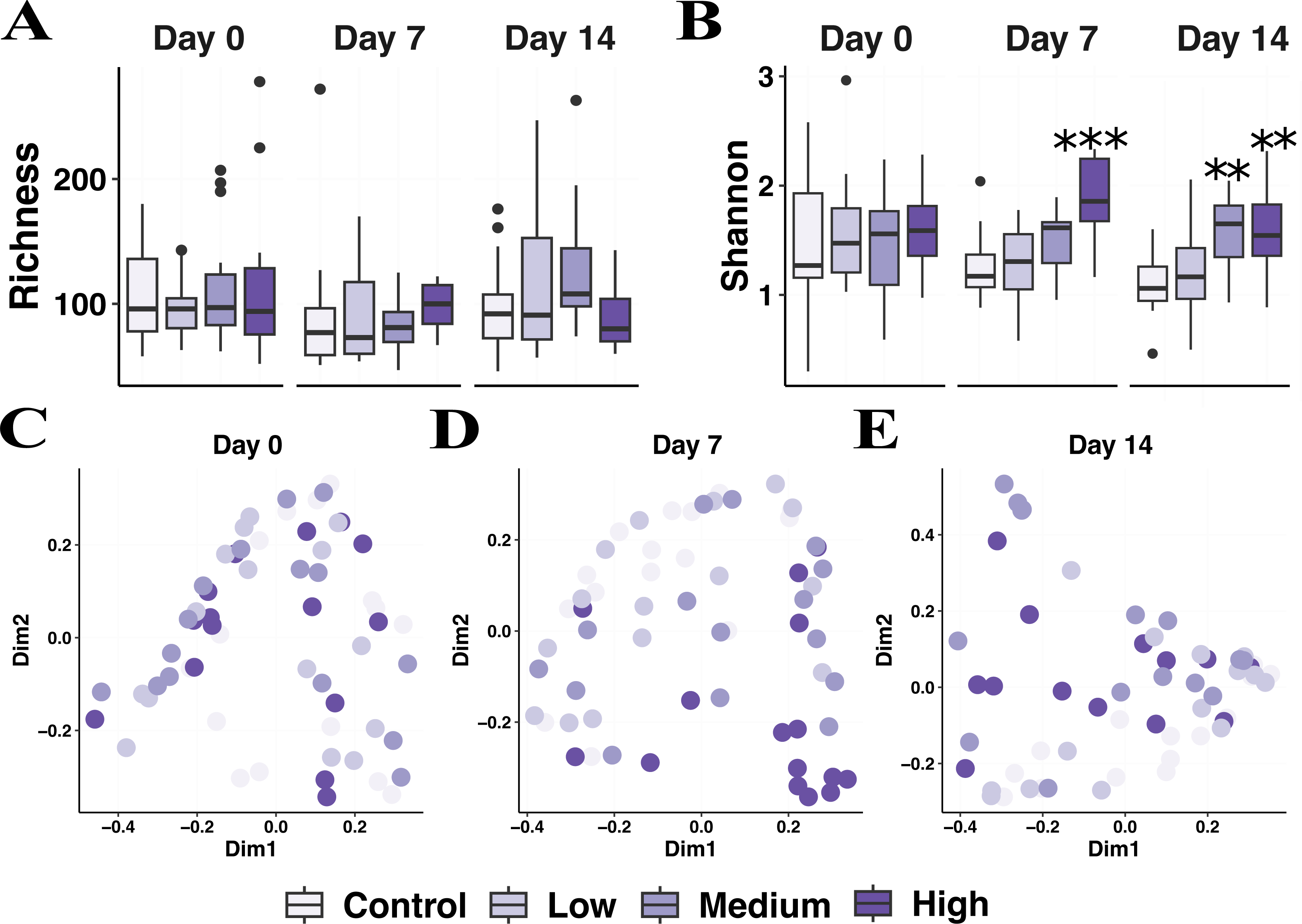
Azoxystrobin exposure disrupts gut microbial diversity and composition. A) Richness and B) Shannon entropy in each exposure group across the length of the experiment. Each box represents the inter quartile range (IQR) of the data, the center line is the median and the whiskers extended 1.5 x IQR beyond the upper and lower hinges. Asterisks indicate significant differences compared to the control group (**, P < 0.01; ***, P < 0.001). Ordination (principal coordinates; Jaccard) of genus-level community profiles at C) day 0, D) day 7, and E) day 14. Each point represents one sample; colors indicate exposure group (n = 15 fish per group; total n = 60).

### Acute AZO exposure alters microbial taxa and functional potential in a sex- and dose-dependent manner

To determine if compositional shifts in microbial communities corresponded to changes in functional potential (*i.e.*, the genes encoded by microbiota) we generated and analyzed shotgun metagenomic libraries from fecal samples collected at Day 7. Day 7 was selected as this was the earliest time point where disruption occurred. Consistent with our 16S rRNA analyses, shotgun metagenomics confirmed that AZO exposure significantly disrupted the gut microbiome on Day 7 (**Figure S2**). Specifically, the number of detected bacterial species (GLMM, z = 2.28, *P* = 2.24 x10^-2^) was significantly increased in high exposed group on day 7 (**Figure S2A)**, and the Shannon diversity of bacteria species was significantly increased in the medium (GLMM, z = 2.45, *P* = 1.44 x10^-2^) and high exposed group (GLMM, z = 5.81, *P* = 6.42 x 10^-9^; **Figure S2B**). The significant effect of the AZO treatment groups on bacterial species composition was observed (PERMANOVA: group R² = 3.33x10^-1^, *P* = 2.00x10^-4^) (**Figure S2C**). At the species level, we confirmed the 16S findings, observing a significant decrease in species from the *Aeromonas* (*A. veronii*), and *Shewanella* (*S. xiamenensis*) genera, while species from *Bosea* (*Bosea sp._ANAM02*) and *Pseudomonas* (*P. mosselii*, *P. aeruginosa* and *P. putida*) genera were increased (**Figure S2D**, **Table S2**).

Given that AZO is a fungicide, we also examined its effects on the gut fungal community. AZO exposure significantly altered fungal diversity and composition on Day 7 (**Figure S2**). Specifically, fungal richness was significantly increased in the medium exposure group (GLMM, z = 2.04, *P* = 4.16 × 10⁻²; **Figure S2E**), and Shannon diversity was significantly increased in both the medium (GLMM, z = 2.49, *P* = 1.28 × 10⁻²) and high exposure groups (GLMM, z = 2.57, *P* = 1.06×10⁻²; **Figure S2F**). In addition, an association between fungal community diversity and treatment group was noted (PERMANOVA: group R² = 2.14x10^-1^, *P* = 4.40×10^⁻3^; **Figure S2G**). Notably, the abundance of the commensal yeast *Saccharomyces cerevisiae* was significantly depleted after high dose of AZO exposure (GLM, z = −1.99, *P* = 4.63 × 10⁻²) (**Figure S2H**).

To gain more insight into the functions that might be disrupted by exposure we asked if exposure resulted in alteration of KEGG pathway abundances. Exposure group, sex, and their interaction collectively explained ∼41% of the variation in zebrafish; the group factor explained the most variation (R^2^ = 0.23, F = 2.58, *P* = 3.98 x10^-2^), followed by the interaction between group and sex (R^2^ = 0.10, F = 1.15, *P* = 3.30 x 10^-1^), and sex (R^2^ = 0.085, F = 2.88, *P* = 7.24 x 10^-2^) (**Figure 2A**), indicating that AZO treatment was one of the major factor driving changes in microbial functional potential. A total of 144 pathways significantly varied across treatment groups (**Figure 2B; Table S3**). Specifically, pathways annotated involved in the degradation of environmental and synthetic compounds were significantly elevated (**Figure 2C**), including those for atrazine degradation (ko00791), xylene degradation (ko00622), chlorocyclohexane and chlorobenzene degradation (ko00361) and aminobenzoate degradation (ko00627). In addition, we observed the abundance significantly increased in pathways associated with the biosynthesis of antimicrobial compounds. Prodigiosin biosynthesis (ko00333), a secondary metabolite with antimicrobial and immunomodulatory properties, was significantly increased in both the medium and high exposure groups. In contrast, several pathways involved in bacterial motility, secretion, and membrane structure were significantly reduced, particularly in high-exposure male zebrafish (**Figure 2C**, **Table S3**). These included flagellar assembly (ko02040) in medium and high exposed groups, bacterial secretion systems (ko03070) in the medium and high exposed groups, and lipopolysaccharide (LPS) biosynthesis (ko00540) in high exposed group.

**Figure 2.**
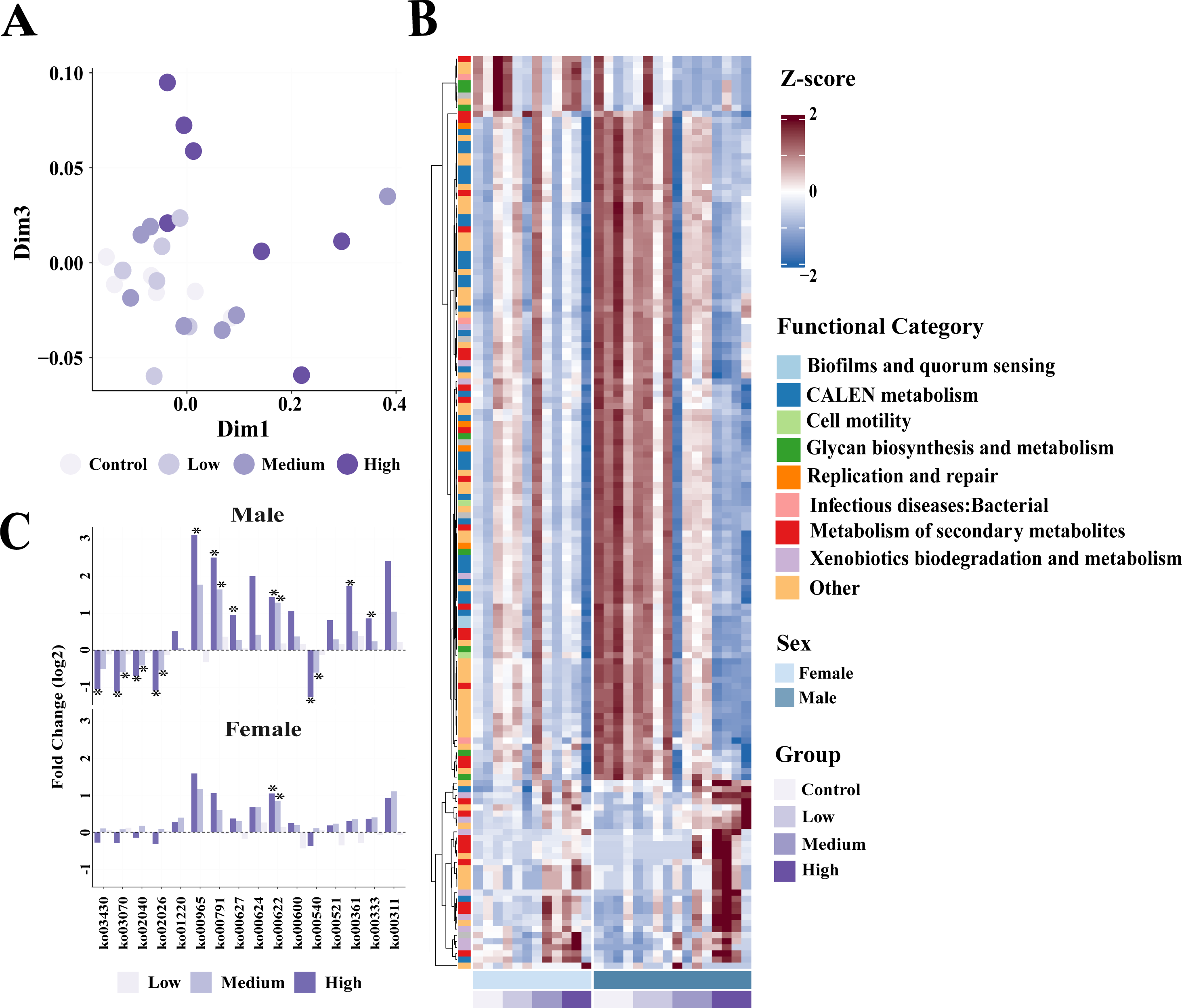
Azoxystrobin exposure rapidly alters gut microbial functional potential. A) An ordination (principal coordinate analysis - PCoA; Jaccard) of KEGG pathway abundances. Group and sex effects were assessed with PERMANOVA (group: R^2^ = 0.23, *P* = 0.039; sex: R^2^ = 0.08, *P* = 0.072). B) Heatmap of significantly varying KEGG pathways (z-score; FDR = 0.2). Each row represents a pathway which varied in abundance between exposed and unexposed groups. Side colors indicate broad KEGG functions. For each column (samples) animal sex is indicated by color bars at the bottom of plot. C) A subset of significantly altered KEGG pathways highlighting alterations in xenobiotic biodegradation and secondary metabolism pathways in males (N = 4/ group) and females (N = 3/group) at day seven.

### Acute AZO exposure induces liver morphological changes and gene expression

After quantifying AZO-induced shifts in the gut microbiome, we evaluated host physiological responses to exposure. We first assessed general body metrics. Although baseline body weight was not recorded, we found after that 14 days of azoxystrobin exposure, body weight (groupLow:sexFemale GLMM, z = −4.32, *P* = 1.58 x10^-5^; groupMedium: sexFemale GLMM, z = −6.06, *P* = 1.32 x10^-9^; groupHigh: sexFemale GLMM, z = −3.62, *P* = 2.95 x10^-4^) and condition factor (groupLow: sexFemale GLMM, z = −3.31, *P* = 9.3 x10^-4^; groupMedium: sexFemale GLMM, z = −4.5, *P* = 6.92 x10^-6^; groupHigh:sexFemale GLMM, z = −3.97, *P* = 7.31 x10^-5^) were significantly lower in high-dose females, while no significant differences in body length were observed across groups (**Figure S3A–C**).

Azoxystrobin (AZO) has been reported to induce hepatic toxicity in toxicology studies (14, 15). Interestingly, there were biliary changes in the livers of fish after 14 days of exposure, with increased severity as dosage increased (**Figure 3**). Control fish exhibited few bile duct profiles per 400x field; ducts were well demarcated, small and surrounded by minimal stromal tissue (**Figure 3A**). In the low-dose group, bile duct profiles were mildly increased and mildly irregular with increased periductal fibrous stroma and ductular reaction extending into the hepatic cords, consistent with a mild degree of biliary hyperplasia (**Figure 3B**). In the medium-dose group, ductular reaction and fibrosis were more extensive (**Figure 3C**), with tortuous bile ducts and ductules, mild lymphocytic inflammation, and occasional karyorhectic cells (**Figure 3D**). High-dose fish exhibited changes similar to those observed in the medium-dose group. Lesion severity was recorded semi-quantitatively (**Table S4**).

**Figure 3.**
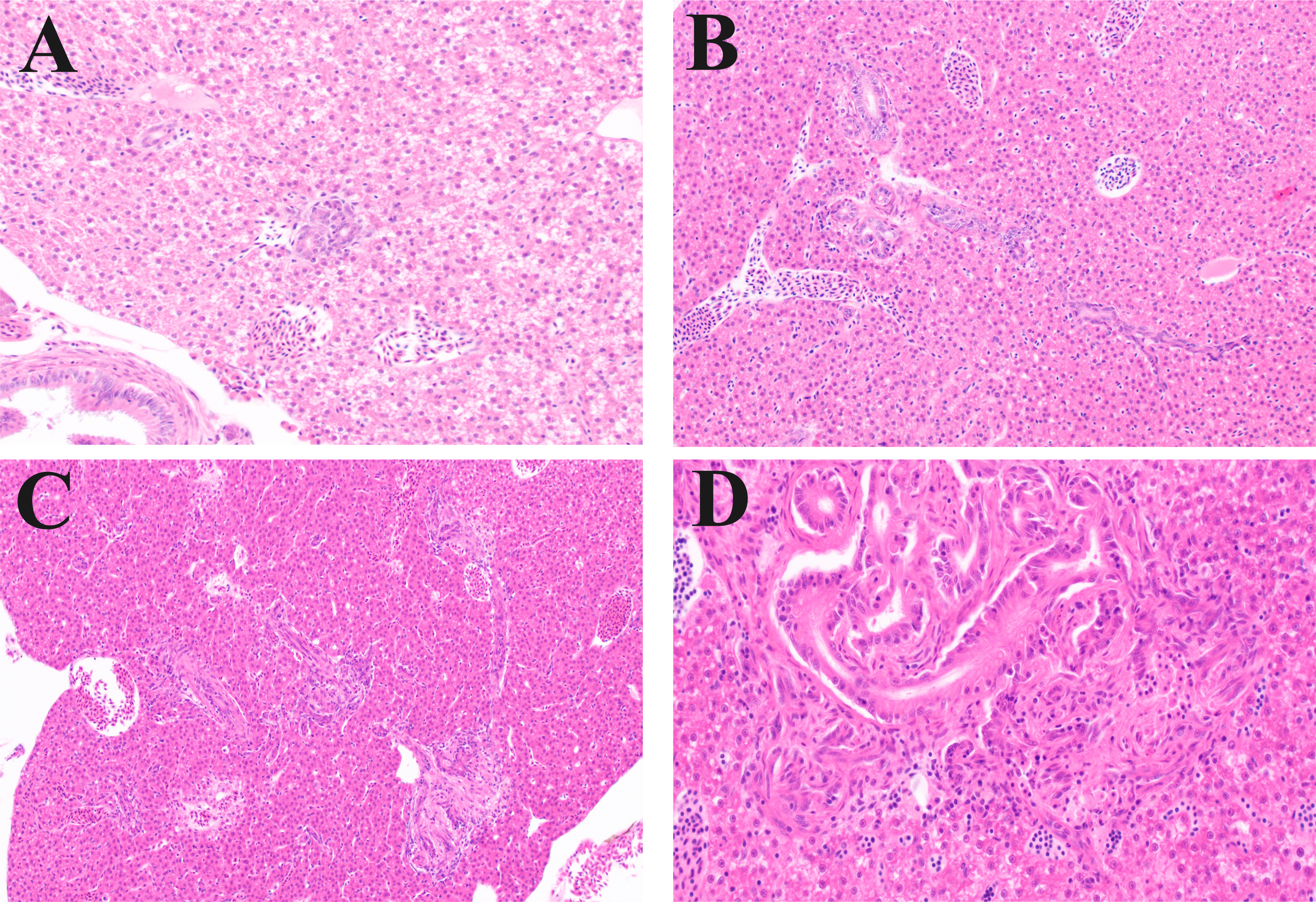
Acute azoxystrobin exposure disrupts hepatic architecture. Representative stained zebrafish liver sections centered on the biliary/periportal region after 14 days of azoxystrobin exposure. A) Control. Bile duct profiles are few within each field, well demarcated, and have minimal stroma. B) Low dose. Bile duct profiles are numerous, with bile ducts lined by ciliated epithelium surrounded by increased fibrous stroma, as well as parenchymal ductular reaction. Considered a mild degree of biliary change. C) Medium dose. Ductular reaction and fibrosis are more extensive. D) Medium dose. Both bile ducts and ductules are tortuous, with mild inflammation (lymphocytes) and few necrotic (karyorrhectic) cells. (200X, A,B,C; 400X D hematoxylin and eosin (H&E) (n = 7 fish per group).

To further investigate impacts of AZO on the liver, we performed transcriptomic profiling of liver tissue from control and high-dose AZO-exposed zebrafish. Differential expression analysis (FDR < 0.1) revealed 143 differentially expressed genes (DEGs) in the combined dataset (52 upregulated, 91 downregulated) (**Figure 4A, Table S5**). Consistent with previous work (14) several genes involved in oxidative stress response and liver injury and repair were altered including *nupr1a* (nuclear protein 1a), reactive oxygen species (ROS) production gene *xdh* (xanthine dehydrogenase), antioxidant and metal-binding protein *mt2* (metallothionein 2), the bile acid nuclear receptor *nr1h4* (FXR, farnesoid x receptor), and the DNA replication factor *mcm7* (minichromosome maintenance complex component 7) were increased. To gain insights into the potential broader biological significance of these DEGs, we performed a Gene Ontology (GO) enrichment analysis (**Table S6;** *FDR* < 0.05). The most significantly upregulated biological processes in exposed zebrafish were related to protein production, including translation (GO:0006412) (*P* = 4.98 × 10⁻^5^) and ribosome biogenesis (GO:0042254) (*P* = 3.11 × 10⁻^4^) (**Figure 4B**). Many downregulated pathways were involved in anabolic metabolism and cellular defense (**Table S6**). Broad functional categories, including lipid metabolic process (GO:0006629) (*P* = 8.09 × 10⁻^5^) and small molecule metabolic process (GO:0044281) (*P* = 2.52 × 10⁻^5^) were significantly enriched for downregulated genes. Key pathways involved in anabolic metabolism and metabolic regulation such as the cholesterol biosynthetic process (GO:0006695) (*P* = 4.48 × 10⁻^5^), acetyl-CoA metabolic process (GO:0006084) (*P* = 2.93 × 10⁻^2^) and insulin receptor signaling pathway (GO:0008286) (*P* = 1.61 × 10⁻^4^) were also enriched for downregulated genes. Consistent with altered oxidative stress response, cell redox homeostasis (GO:0045454) (*P* = 3.28 × 10⁻^2^) was enriched for downregulated genes, including decreased expression of *gsr* (glutathione-disulfide reductase) and *prdx1* (peroxiredoxin 1).

**Figure 4.**
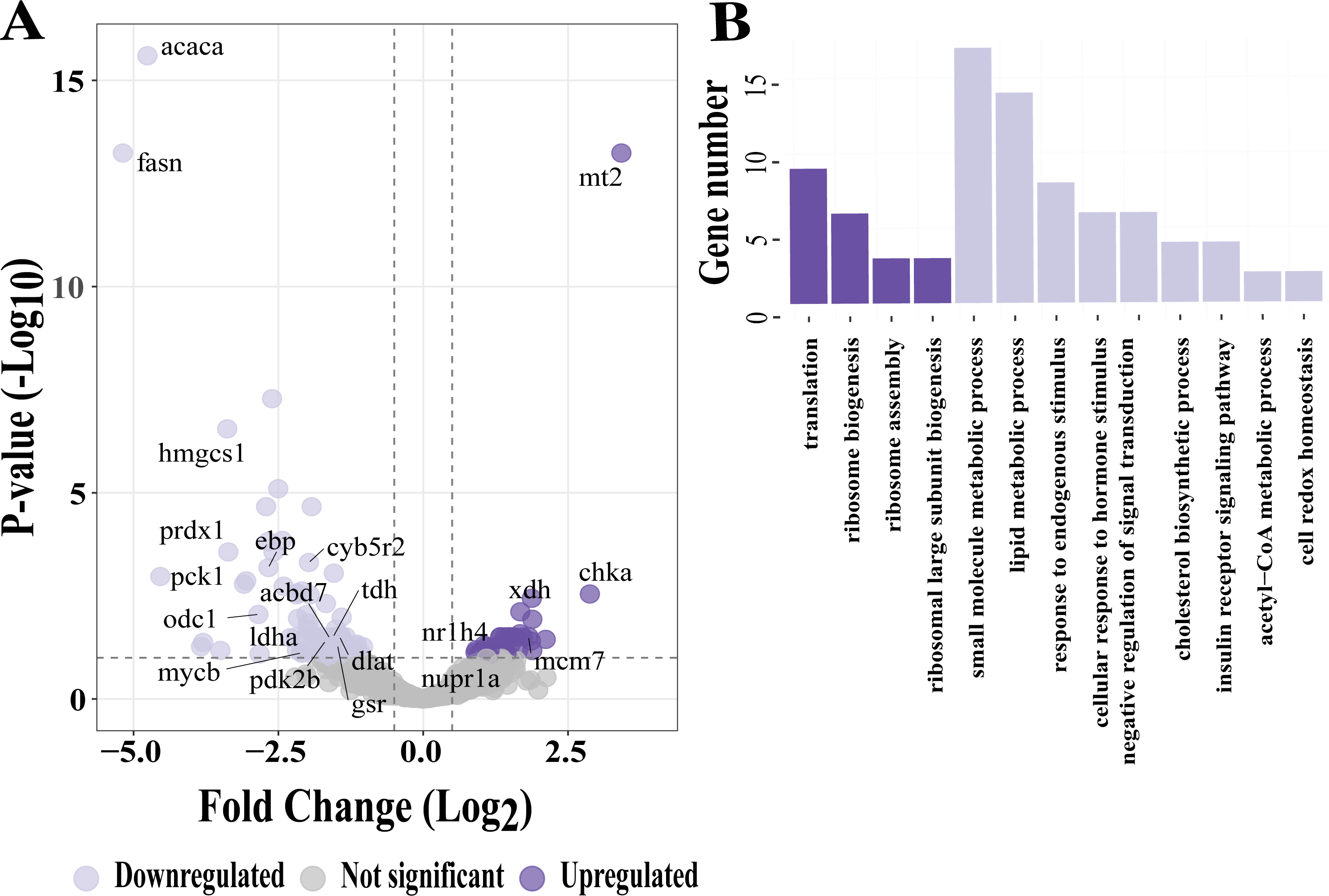
Azoxystrobin exposure disrupts hepatic gene expression. A) Volcano plot depicting differentially expressed genes in high-exposure group relative to control animals (control, n = 7; high, n = 8; FDR < 0.10). B) Bar plot of biological processes enriched for upregulated (dark purple) and downregulated (light purple) gene (FDR < 0.05).

Consistent with prior reports of sex specific toxicant response, hepatic transcriptional response to AZO was found to be highly sexually dimorphic. Specifically, in females we identified 76 upregulated and 95 downregulated genes (**Figure S4A**, **Table S7**) while 25 genes increased and 85 decreased in males (**Figure S4B, Table S8**). Few genes were altered in the same way across sexes, but both males and female downregulated genes were commonly enriched in small molecule metabolism and cholesterol biosynthesis (**Figure S4C, D**). Male down regulated genes are also enriched in metabolic pathways such as gluconeogenesis (**Table S9**) and lipid metabolism while genes were enriched in cell redox homeostasis in females (**Table S10**).

### Azoxystrobin-induced changes in microbiota correlate with hepatic stress and metabolic gene expression in zebrafish

To explore the relationship between microbiota composition and hepatic transcriptional responses under azoxystrobin (AZO) exposure, we conducted Spearman correlation analyses between significantly altered bacterial genera (16S rRNA) and differentially expressed liver genes (DEGs). We find that 2,431 unique associations between 17 taxa and 143 genes (**Figure 5A**). More specifically, several genera were strongly correlated with genes involved in oxidative stress and metabolic regulation (**Figure 5B**). Across all fish, increased relative abundance of *Pseudomonas*, *Shewanella*, *Bosea*, *Mycoplana*, and *Paenirhodobacter* was significantly associated with elevated expression of translation related genes (*rpl37*, *rps11*, *rpl9*, *rps15*, *rpl35*, *rps17*, *rpl35a*, *rpl13*, *rps14*) and oxidative stress related genes such as *mt2*, *xdh*, and *nupr1a*, and *nr1h4* (FXR), a central regulator of bile acid homeostasis, consistent with the biliary-associated histopathological changes observed under AZO exposure (FDR < 0.15; **Figure 5B**). In contrast, the decreased abundance of genera such as *Phreatobacter*, *Weissella*, *Leuconostoc*, and *Vagococcus* showed positive correlations with genes involved in small molecule metabolic processes such as *fasn*, *acaca*, *apof*, and *hmgcs1*(**Figure 5B**).

**Figure 5.**
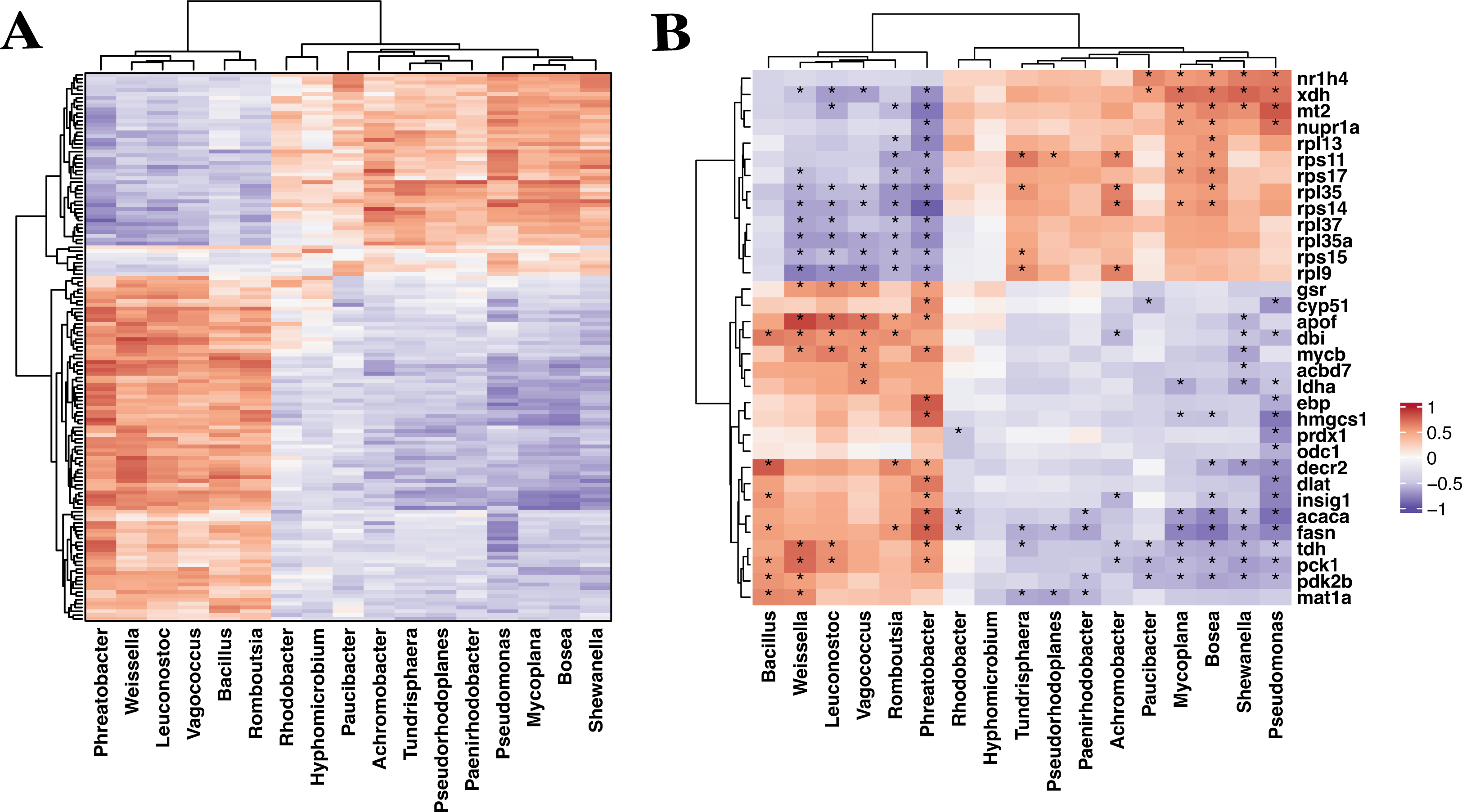
Disruption of gut microbial communities correlates with altered liver gene expression. **A)** Spearman correlation heatmap between the relative abundance of significantly altered 16S rRNA genera and liver differentially expressed genes (DEGs) (control, n = 7; high, n = 8). **B)** Heatmap highlighting correlations between genera and genes in the small-molecule metabolic process gene set (downregulated in the high-exposure group) and the translation-related gene set (upregulated in the high-exposure group). In both panels, colors indicate Spearman’s correlation coefficients (red, positive; purple, negative). Asterisks denote significant correlations with |ρ| > 0.3 and FDR < 0.15. Row and column cladograms indicate hierarchically clustered to reveal co-association patterns.

When stratified by sex, distinct microbiota–host expression relationships emerged. In females 688 associations were identified between 4 taxa and 172 genes. Significant positive correlations were observed between *Paucibacter* and *Pseudomonas* and genes involved in cell cycle regulation (*ccna2*, *cdc6*, *ncapd2* and *anln*) and oxidative stress related genes (*mt2* and *nupr1b*), while *Phreatobacter* and *Alistipes* were negatively associated with key metabolic genes such as *fasn*, *hmgcs1*, *cyp51*, and *acaca* (**Figure S5A–B**). In males, 1980 associations were identified between 18 taxa and 110 genes. Strong positive correlations were observed between *Shewanella*, *Mycoplana*, *Pseudomonas*, and *Reyranella*, and the oxidative stress marker *xdh*, *mt2*, *stat1a* and *bbox1* (**Figure S5C–D**). Several of these same genera were also negatively correlated with lipid metabolic genes, including *fasn*, *acaca*, *pck1*, and *hmgcs1*. Conversely, the decreased abundance of genera such as *Weissella*, *Leuconostoc*, *Vagococcus*, and *Peptostreptococcus* were positively correlated with the genes involving in lipid metabolic genes, including *fasn*, *acaca*, *pck1*, and *hmgcs1*. Together, these findings reveal coordinated changes in microbial composition and hepatic gene expression in response to AZO exposure.

## Discussion

Azoxystrobin (AZO) is one of the most extensively applied fungicides worldwide (1, 2), and residues have been detected in food (1), water (7), and humans, including pregnant women and children (9). Despite its widespread use, how AZO exposure shapes the gut microbiome and how these changes relate to downstream host physiology remains incompletely understood. This matters because the gut microbiome is increasingly recognized as an important determinant of xenobiotic metabolism and host metabolic signaling with relevance to hepatic function (17, 18). Here, we addressed this gap by integrating microbial and hepatic endpoints to show that AZO exposure is associated with dose and sex-dependent disruptions in microbial communities and liver histopathology and gene expression. While more reductionist experimentation is needed to evaluate directionality and specific mechanisms, our findings suggest that AZO associated disruption of microbial community composition and genetic repertoire may contribute to exposure impacts in the liver.

Our study demonstrates that acute AZO exposure significantly restructures gut microbiome composition and function. Prior studies have shown that AZO can perturb gut microbial communities in other host systems (16, 38). Here we extend these observations by showing that microbial functional potential also shifts rapidly in response to AZO exposure in zebrafish. This is reflected in the increased abundance of multiple pathways involved in xenobiotic metabolism. Notably, the abundance of pathways for atrazine, xylene, chlorobenzene, and aminobenzoate degradation were significantly increased despite these chemicals not being present. This is plausibly related to the chemical structure of AZO, which contains chlorophenyl, cyanophenyl, and aromatic moieties, and may reflect engagement of broad-spectrum microbial enzymes such as oxygenases, dehalogenases, and amidases (39, 40). Together, these shifts are consistent with selective pressures that could favor taxa with enzymatic capacity to transform AZO or related compounds. This interpretation is supported by recent mechanistic studies reporting genes and enzymes capable of azoxystrobin degradation. For instance, Jiang et al. cloned and characterized a novel esterase gene, *strh*, from a *Hyphomicrobium* species that can hydrolyze strobilurin fungicides (41). The enzyme encoded by this gene, *StrH*, hydrolyzes the ester bond of azoxystrobin to form its primary, less toxic metabolite, azoxystrobin acid (41). It is possible that AZO disruption of gut microbial communities free niche space for microbes that possess the metabolic capacity to degrade AZO thereby limiting its harms. Alternatively, incoming microbes may possess metabolic pathways that generate metabolic endpoints or intermediates that, while benign to the microbe encoding them, may be more toxic to other microbes or contribute to hepatic stress responses and metabolic disruption of the host (42). This scenario is consistent with the transcriptional disruption of lipid metabolism and patterns we observed. More work is needed to determine if direct microbial metabolism of AZO mitigates or contributes directly to exposure associated pathologies.

Exposed animals possessed lower abundance of microbial pathways involved in flagellar motility, secretion systems, and lipopolysaccharide (LPS) biosynthesis. These functions can shape colonization dynamics and host sensing at the host-microbe interface (43). For example, LPS influences host immunity, gut homeostasis, inflammation and can help resist immune responses (44). A simple possible explanation for decreases in LPS genes is a compositional shift towards a more gram-positive rich microbiota in exposed animals. However, we found few gram-positive organisms were significantly increased during exposure although several gram-negatives decreased. Regardless of the cause the observed reduction in LPS gene abundance suggests that AZO exposure may alter signaling through microbial associated molecular patterns which could in turn could impact host immune function and local inflammation (45). Concurrent with these shifts, the liver exhibited transcriptomic evidence of disrupted redox balance, impaired lipid metabolism, and loss of regulatory pathways. This cooccurrence raises the possibility that the altered microbial signaling may alter hepatic responses to xenobiotic stress (46).

An important finding of this work is that these aspects of shifts in microbial communities are sex-specific and co-occur with sexually dimorphic hepatic responses. Sex-dependent susceptibility to pesticides has been well described (47). Here, we show that microbiome and liver responses together exhibit sex-stratified patterns, consistent with sex as an important source of variation. Specifically, male zebrafish showed significant alteration in more pathways associated with xenobiotic metabolism and a hepatic transcriptomic signature suggestive of oxidative-stress responses and metabolic disruption. In contrast, females exhibited microbiota–host expression associations that more strongly involved cell-cycle– related genes, alongside a hepatic transcriptional profile consistent with cellular remodeling and repair. These findings are consistent with reports that sex hormones shape the gut microbiome and influence systemic outcomes (48, 49) and suggest that the microbiome may be a contributor to sex-specific toxicological responses. Sex-stratified responses may also reflect sex differences in internal exposure (toxicokinetics) and baseline host physiology (47, 50). Together, these results underscore the importance of considering sex as a biological variable in microbiome-toxicant studies.

Liver histopathology identified biliary hyperplasia with ductular reaction, mild inflammation, and mild biliary cell necrosis after AZO exposure, evident in medium-dose and high-dose fish. Ductular reactions are commonly observed in cases of cholestasis or biliary injury (51). At the molecular level, AZO exposure was also accompanied by hepatic transcriptional changes consistent with a stress response, including altered redox-related genes and reduced expression of genes involved in lipid and small-molecule metabolism. Notably, expression of the bile acid nuclear receptor FXR (*nr1h4*) was significantly increased during exposure, which may reflect altered bile acid signaling or a compensatory response to periportal/biliary perturbation during exposure. Because bulk RNA-seq averages gene expression across mixed cell populations and anatomical regions, localized periportal and biliary changes can be difficult to resolve in whole-liver transcriptomes, motivating spatially resolved approaches in future studies (51–54). Taken together, these endpoints describe concurrent histologic and transcriptional responses during acute exposure, while the upstream drivers and temporal sequence remain to be established.

To elucidate potential mechanistic links between the observed microbial shifts and hepatic endpoints, we performed correlation analysis to identify candidate drivers along the gut-liver axis. Notably, the specific expansion of opportunistic *Proteobacteria*, including *Pseudomonas* and *Shewanella*, was strongly correlated with the upregulation of the bile acid regulator *nr1h4* (FXR) and hepatic oxidative stress genes (*mt2*, *xdh*). This molecular association parallels the observed histopathological biliary reaction. Since *Pseudomonas* species possess the enzymatic capacity to metabolize bile acids (55), which serve as endogenous ligands for FXR, their expansion could potentially alter the bile acid pool, thereby contributing to the observed dysregulation of FXR signaling and biliary homeostasis (56). Furthermore, members of the *Pseudomonas* genus are well-documented producers of redox-active factors, such as pyocyanin, capable of generating reactive oxygen species (57). The expansion of these redox-active taxa offers a plausible explanation for the observed upregulation of *mt2*, which functions as one of the primary hepatic defenses against microbial-derived oxidative stress (58). Conversely, the depletion of lactic acid bacteria, such as *Weissella* and *Leuconostoc*, was positively correlated with the downregulation of lipid synthesis genes (*fasn*, *acaca*). As these genera are key producers of lipogenic substrates like acetate, their loss offers a potential mechanistic basis for the suppressed hepatic lipid metabolism observed in the transcriptomic profiles (59). Collectively, these associations suggest that AZO-induced hepatotoxicity may be compounded by a dual mechanism: the metabolic disruption of bile and stress signaling by expanding opportunists and the loss of substrate support from commensals.

The microbiome-host interactions observed here align with observations reported in pesticide toxicology, suggesting a potential signature of chemically induced dysbiosis. The expansion of *Proteobacteria* is frequently attributed to the selection of taxa capable of xenobiotic resistance or degradation (60). Notably, the linkage between dysbiosis and suppressed hepatic lipogenesis resembles responses reported for other fungicide and insecticide exposures. For instance, Wang et al. and Jin et al. reported that exposure to chlorpyrifos or propamocarb in zebrafish was associated with reduced expression of hepatic lipogenic genes (e.g., *fasn*, *acc1*), coinciding with *Proteobacteria* expansion and oxidative-stress signatures (61, 62). Together, these parallels suggest that AZO-associated suppression of host lipid metabolism may be part of a shared response pattern to pesticide stress involving the gut–liver axis.

Together our data suggest that acute exposure to AZO disrupts microbial composition and functional potential and that these changes associate with altered liver physiology. One caveat of this work as that our results are largely correlative and future mechanistic study is needed to establish a causative role for the microbiome in these responses. This work is also limited in that it only examines acute timeframes of exposure. It is possible that longer exposure periods would yield effects even at lower exposure concentrations. As many environmental exposures occur over long timeframes at low doses (e.g., lead exposure) it will be important to clarify how exposure duration impact host-microbiota toxicant interactions. Despite these limitations, our findings support the growing body of literature that suggests that the microbiome plays an important role in modulating the impact of environmental exposures on host health raising the possibility that microbiome-based therapeutics may one day be used to modulate or prevent exposure harms.

## Acknowledgements

This work was supported by the National Institute of Environmental Health Sciences of the National Institutes of Health under Award Number R01ES036174 to CAG. The content is solely the responsibility of the authors and does not necessarily represent the official views of the National Institutes of Health. This work was also supported by institutional funds to CAG. LD was partially supported by Illinois Distinguished Fellowship for a graduate student at the University of Illinois at Urbana-Champaign (UIUC). Authors thank the Roy J. Carver Biotechnology Center for DNA sequencing service and the UIUC College of Veterinary Medicine Shared Equipment Program and the Biocomputing Shared Resource (BioShaRe) for use of their equipment. This study made use of the Illinois Campus Cluster, a computing resource that is operated by the Illinois Campus Cluster Program (ICCP) in conjunction with the National Center for Supercomputing Applications (NCSA), which is supported by funds from the UIUC.

